# Reverse transcriptase fused CRISPR-Cas1 locus with RNA-seq expression necessitates revisiting hypothesis on acquisition of antibiotic resistance genes in multidrug-resistant *Enterococcus faecalis* V583

**DOI:** 10.1101/263558

**Authors:** Sandeep Chakraborty

## Abstract

The emergence of drug-resistance in *Enterococcus faecalis* V583 through acquisition of resistance genes has been correlated to the absence of CRISPR-loci. Here, the presence of a bona-fide CRISPR locus in E. faecalis V583 (Accid:NC_004668.1) at 2238156 with a single 20 nt repeat is demonstrated. The presence of a putative endonuclease Cas1 13538 nucleotides away from the repeat substantiates this claim. This Cas1 (628 aa) is highly homologous (Eval:5e-34) to a Cas1 from *Pseudanabaena biceps* (Accid:WP 009625648.1, 697 aa), which belongs to the enigmatic family of RT-CRISPR locus. Such significant similarity to a Cas protein, the presence of a topoisomerase, other DUF (domain of unknown function) proteins as is often seen in CRISPR loci, and other hypothetical proteins indicates that this is a bona-fide CRISPR locus. Further corroboration is provided by expression of both the repeat and the Cas1 gene in existing RNA-seq data (SRX3438611). Since so little is known of even well-studied species like E. *faecalis* V583 having many hypothetical proteins, computational absence of evidence should not be taken as evidence of absence (both crisprfinder and PILER-CR do not report this as a CRISPR locus). It is unlikely that bacteria would completely give up defense against its primeval enemies (viruses) to bolster its fight against the newly introduced antibiotics.

## Introduction

Antibiotic-resistant bacterial infections are linked to increased morbidity and mortality in hospitals [1]. *Enterococcus faecalis* is a gram-positive, commensal bacteria found in the gastrointestinal tract of humans.One strain, *E. faecalis V583,* has developed resistance to vancomycin [2] and chloramphenicol treatment causing great concern [3]. Mobile DNA, easily transferred among different strains and species, has been shown to be a large contributing factor to this resistance [4, 5].

Clustered regularly interspaced short palindromic repeats (CRISPR) is a prokaryotic adaptive defense system in bacteria [6]. Through this mechanism bacteria assimilate (memorize) short sequences of invading genomes (spacers) within repeats [7–9], and uses proximal effector proteins (Cas) [10–12] to cleave subsequent infections from the same organism [13, 14]. Although CRISPR-Cas loci have been identified in some *E faecalis* strains (for e.g. T1 and X98) [15–17], it was not found in *E. faecalis V583* [16]. Subsequent work has correlated the emergence of drug-resistance in *E. faecalis* V583 to the absence of CRISPR-loci [16, 18, 19].

Here, a CRISRP-Cas locus is reported in *E. faecalis V583,* with expression found for both the repeat and the Cas1 gene in existing RNA-seq data.

## Results and discussion

### Repeat and proximal Cas1 gene

The 20 nt repeat at 2238156 (GGGCAGTCGGCGTTGACCGC) is shown in Fig 1. The presence of an putative endonuclease Cas1 (628 aa) 13538 nucleotides away from the repeat substantiates this claim (Table 1). This Cas1 is highly homologous (Eval:5e-34) to the N-terminal of a Cas1 from *Pseudanabaena biceps* (Accid:WP_009625648.1, 697 aa), which is annotated as a group II intron reverse transcriptase (RT). The C-terminal of this protein has been established to have nucleolytic activity using a homolog from *Archaeoglobus Fulgidus* (PDBid:4N06), indicating this belongs to the enigmatic RT-CRISPR family [20–22].

### RNA-seq evidence

Expression was found for both the repeat (SRR6339668.9502786.1, SRR6339668.8954887.2, SRR6339668.4564129.1, SRR6339668.4376029.2, SRR6339668.1205782.2) and the Cas1 gene (Fig 2) based on the SRA:SRX3438611.

### Conclusions

Computational absence of evidence should not be taken as evidence of absence. Neither crisprfinder [23], nor PILER-CR [24] report this as a CRISPR locus. Repeats with relevant proximal proteins should give enough credence to look for CRISPR-loci [25]. Claims that ‘major bacterial lineages are essentially devoid of CRISPR-Cas viral defense systems’ should also be revisited [26], since its unlikely they have failed to “learn” in billions of years the CRISPR defense mechanism against viruses from their peers - something that they did in decades against human introduced antibiotics. For example, the claim that the entire *Chlamydiae* phylum lack CRISPR-Cas systems [26] was shown to be mistaken [27]. Also, it is unlikely that bacteria would give up defense against its primeval enemies (viruses) to bolster its fight against the newly introduced antibiotics.

## Materials and methods

ftp://ftp.ncbi.nih.gov/genomes/refseq/bacteria/ provided bacterial genomes and annotation. The getorf program from the EMBOSS suite [13] was used to obtain the ORFs (SI ORFS.fa). ORFs that are 15000 nt from the loci are annotated on both sides using YeATS [28-30] based on the Ncbi annotation. MSA figures were generated using the ENDscript server [31]. RNAfold [32] was used to predict the secondary structure of the RNA repeat. MAFFT (v7.123b) [33] was used for doing the multiple sequence alignment (MSA).

## Competing interests

No competing interests were disclosed.

**Table 1.** Open reading frames (ORF) within 15000 nucleotides of a repeat at 2238156: These are BLAST results, except for the asterisked line, which shows that the ORF NC_004668.1_1982 (628 aa) is homologous to the Cas1 protein (Accid:WP_009625648.1) from *Pseudanabaena biceps,* where the N-terminal is annotated as a reverse transcriptase and the C-terminal has nucleolytic activity. Several hypothetical proteins and CRISPR relevant genes like topoisomerases and DUFs are also located in this region. Finally, RNA-seq data shows expression of both the repeat and the Cas1 gene. See SI ORFS.fa for sequences and location.

| Ordered ORF | Length | Accid | Description | BBS | Evalue |
| --- | --- | --- | --- | --- | --- |
| NC_004668.1_1981 | 555 | WP_002368726.1 | MULTISPECIES: conjugal transfer protein TraG | 1231 | 0 |
| NC_004668.1_1982 | 628 | WP_011109539.1 | group II intron reverse transcriptase/maturase | 1308 | 0 |
| *NC_004668.1_1982 | 628 | WP_009625648.1 | **CRISPR-associated endonuclease Cas1** | 135 | 5e-34 |
| NC_004668.1_1983 | 289 | WP_002368724.1 | MULTISPECIES: hypothetical protein | 585 | 0 |
| NC_004668.1_1984 | 184 | WP_002368723.1 | MULTISPECIES: DNA methyltransferase | 352 | 2e-122 |
| NC_004668.1_1985 | 434 | WP_010773628.1 | MULTISPECIES: DUF3848 domain-containing protein | 893 | 0 |
| NC_004668.1_1986 | 139 | WP_002368721.1 | MULTISPECIES: PrgI family protein | 285 | 3e-97 |
| NC_004668.1_1987 | 753 | WP_010773627.1 | MULTISPECIES: conjugal transfer protein TraE | 1707 | 0 |
| NC_004668.1_1988 | 364 | WP_010773626.1 | MULTISPECIES: hypothetical protein | 759 | 0 |
| NC_004668.1_1989 | 910 | WP_010773625.1 | MULTISPECIES: peptidase M24 | 1916 | 0 |
| NC_004668.1_1990 | 290 | EEU68006.1 | radical SAM domain-containing protein | 622 | 0 |
| NC_004668.1_1991 | 746 | WP_002368715.1 | DUF4366 domain-containing protein | 1493 | 0 |
|  |  | Repeat at 2238156 |  |  |  |
| NC_004668.1_1994 | 705 | WP_002368714.1 | MULTISPECIES: DNA topoisomerase III | 1448 | 0 |
| NC_004668.1_1995 | 196 | EEU68001.1 | predicted protein | 392 | 8e-138 |
| NC_004668.1_1998 | 3173 | WP_010774229.1 | MULTISPECIES: hypothetical protein | 6569 | 0 |

**Figure 1:**
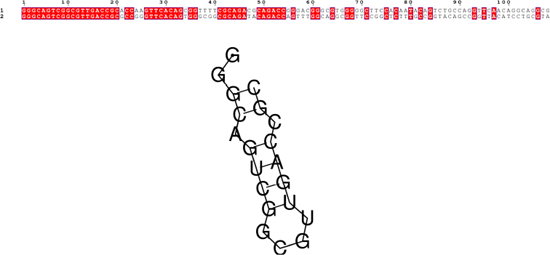
Repeats and spacers in the new locus: The sequence GGGCAGTCGGCGTTGACCGC (20 nt) is repeated once at 2238156, and encompasses a fragment 88 nucleotides long. The secondary structure of the crna is predicted by RNAfold [32]. Note there two mismatched pairs. The cleavage site is predicted to the sequence at the end of the stem on the 3’ end [34].

**Figure 2:**
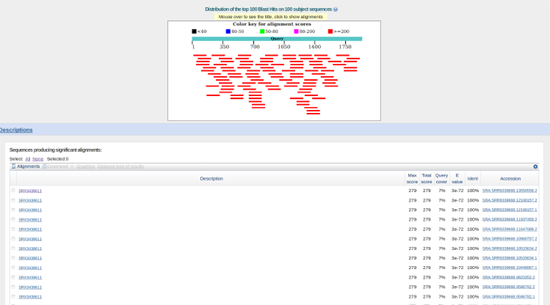
Transcripts of the putatative Casi gene in RNA-seq (SRA:SRR633966) Transcripts matching to the ORF encoded by 2249862 to 2251802 (NC_004668.1_1982, see SI ORFS.fa) - see Table 1.

## References

1. Wisplinghoff H, Bischoff T, Tallent SM, Seifert H, Wenzel RP, et la. (2004) Nosocomial bloodstream infections in us hospitals: analysis of 24,179 cases from a prospective nationwide surveillance study. Clinical infectious diseases 39: 309–317.

2. Arias CA, Murray BE (2012) The rise of the enterococcus: beyond vancomycin resistance. Nature Reviews Microbiology 10: 266.

3. Aakra Å, Vebø H, Indahl U, Snipen L, Gjerstad Ø, et la. (2010) The response of enterococcus faecalis v583 to chloramphenicol treatment. International Journal of Microbiology 2010.

4. Paulsen IT, Banerjei L, Myers G, Nelson K, Seshadri R, et la. (2003) Role of mobile dna in the evolution of vancomycin-resistant enterococcus faecalis. Science 299: 2071–2074.

5. Noble W, Virani Z, Cree RG (1992) Co-transfer of vancomycin and other resistance genes from enterococcus faecalis nctc 12201 to staphylococcus aureus. FEMS microbiology letters 93: 195–198.

6. Ishino Y, Shinagawa H, Makino K, Amemura M, Nakata A (1987) Nucleotide sequence of the iap gene, responsible for alkaline phosphatase isozyme conversion in escherichia coli, and identification of the gene product. Journal of bacteriology 169: 5429–5433.

7. Mojica FJ, García-Martínez J, Soria E, et la. (2005) Intervening sequences of regularly spaced prokaryotic repeats derive from foreign genetic elements. Journal of molecular evolution 60: 174–182.

8. Bolotin A, Quinquis B, Sorokin A, Ehrlich SD (2005) Clustered regularly interspaced short palindrome repeats (crisprs) have spacers of extrachromosomal origin. Microbiology 151: 2551–2561.

9. Horvath P, Barrangou R (2010) Crispr/cas, the immune system of bacteria and archaea. Science 327: 167–170.

10. Barrangou R, Fremaux C, Deveau H, Richards M, Boyaval P, et la. (2007) Crispr provides acquired resistance against viruses in prokaryotes. Science 315: 1709–1712.

11. Jinek M, Chylinski K, Fonfara I, Hauer M, Doudna JA, et la. (2012) A programmable dual-rna-guided dna endonuclease in adaptive bacterial immunity. Science 337: 816–821.

12. Mali P, Yang L, Esvelt KM, Aach J, Guell M, et la. (2013) Rna-guided human genome engineering via cas9. Science 339: 823–826.

13. Marraffini LA (2015) Crispr-cas immunity in prokaryotes. Nature 526: 55–61.

14. Sander JD, Joung JK (2014) Crispr-cas systems for editing, regulating and targeting genomes. Nature biotechnology 32: 347–355.

15. Hullahalli K, Rodrigues M, Schmidt BD, Li X, Bhardwaj P, et la. (2015) Comparative analysis of the orphan crispr2 locus in 242 enterococcus faecalis strains. PloS one 10: e0138890.

16. Palmer KL, Gilmore MS (2010) Multidrug-resistant enterococci lack crispr-cas. MBio 1: e00227–10.

17. Horvath P, Coûte-Monvoisin AC, Romero DA, Boyaval P, Fremaux C, et la. (2009) Comparative analysis of crispr loci in lactic acid bacteria genomes. International journal of food microbiology 131: 62–70.

18. Hullahalli K, Rodrigues M, Palmer KL (2017) Exploiting crispr-cas to manipulate enterococcus faecalis populations. Elife 6.

19. Hullahalli K, Rodrigues M, Nguyen U, Palmer K (2017) A semi-lethal crispr-cas system permits dna acquisition in enterococcus faecalis. bioRxiv : 232322.

20. Toro N, Martínez-Abarca F, González-Delgado A (2017) The reverse transcriptases associated with crispr-cas systems. Scientific reports 7: 7089.

21. Silas S, Mohr G, Sidote DJ, Markham LM, Sanchez-Amat A, et la. (2016) Direct crispr spacer acquisition from rna by a natural reverse transcriptase-cas1 fusion protein. Science 351: aad4234.

22. Silas S, Makarova KS, Shmakov S, Páez-Espino D, Mohr G, et la. (2017) On the origin of reverse transcriptase-using crispr-cas systems and their hyperdiverse, enigmatic spacer repertoires. MBio 8: e00897–17.

23. Grissa I, Vergnaud G, Pourcel C (2007) Crisprfinder: a web tool to identify clustered regularly interspaced short palindromic repeats. Nucleic acids research 35: W52–W57.

24. Edgar RC (2007) Piler-cr: fast and accurate identification of crispr repeats. BMC bioinformatics 8: 18.

25. Chakraborty S (2018) An atypical crispr-cas locus in symbiobacterium thermophilum flanked by a transposase, a reverse transcriptase, the endonuclease muts2 and a putative cas9-like protein. bioRxiv : 252296.

26. Burstein D, Sun CL, Brown CT, Sharon I, Anantharaman K, et la. (2016) Major bacterial lineages are essentially devoid of crispr-cas viral defence systems. Nature communications 7: 10613.

27. Bertelli C, Cissé OH, Rusconi B, Kebbi-Beghdadi C, Croxatto A, et la. (2016) Crispr system acquisition and evolution of an obligate intracellular chlamydia-related bacterium. Genome biology and evolution 8: 2376–2386.

28. Chakraborty S, Britton M, Wegrzyn J, Butterfield T, Martinez-Garcia PJ, et la. (2015). YeATS-a tool suite for analyzing RNA-seq derived transcriptome identifies a highly transcribed putative extensin in heartwood/sapwood transition zone in black walnut.

29. Chakraborty S, Britton M, Martínez-García P, Dandekar AM (2016) Deep RNA-seq profile reveals biodiversity, plant-microbe interactions and a large family of NBS-LRR resistance genes in walnut (juglans regia) tissues. AMB Express 6: 1.

30. Martínez-García PJ, Crepeau MW, Puiu D, Gonzalez-Ibeas D, Whalen J, et la. (2016) The walnut (juglans regia) genome sequence reveals diversity in genes coding for the biosynthesis of nonstructural polyphenols. The Plant Journal.

31. Robert X, Gouet P (2014) Deciphering key features in protein structures with the new endscript server. Nucleic acids research 42: W320–W324.

32. Mathews DH, Disney MD, Childs JL, Schroeder SJ, Zuker M, et la. (2004) Incorporating chemical modification constraints into a dynamic programming algorithm for prediction of rna secondary structure. Proceedings of the National Academy of Sciences of the United States of America 101: 7287–7292.

33. Katoh K, Standley DM (2013) MAFFT multiple sequence alignment software versión 7: improvements in performance and usability. Mol Biol Evol 30: 772–780.

34. Sashital DG, Jinek M, Doudna JA (2011) An rna-induced conformational change required for crispr rna cleavage by the endoribonuclease cse3. Nature structural & molecular biology 18: 680–687.

